# Genistein enhances NAD^+^ biosynthesis by binding to Prohibitin 1 and upregulating nicotinamide phosphoribosyltransferase in adipocytes

**DOI:** 10.1101/2023.04.20.537596

**Authors:** Shun Watanabe, Riki Haruyama, Koji Umezawa, Ikuo Tomioka, Soichiro Nakamura, Shigeru Katayama, Takakazu Mitani

## Abstract

Decreased NAD^+^ levels in adipocytes cause adipose-tissue dysfunction, leading to systemic glucose and lipid metabolism failure. Therefore, developing small molecules and nutraceuticals that can increase NAD^+^ levels in adipocytes is necessary. Genistein, a nutraceutical derived from soybeans, has various physiological activities and improves glucose and lipid metabolism. In this study, we aimed to unravel the effects of genistein on the intracellular NAD^+^ levels in adipocytes and the underlying molecular mechanisms. We showed that genistein enhanced NAD^+^ biosynthesis by increasing the expression of nicotinamide phosphoribosyltransferase (NAMPT), the rate-limiting enzyme in NAD^+^ biosynthesis. A pull-down assay using genistein-immobilized beads identified prohibitin 1 (PHB1) as a target protein of genistein. The knockdown of PHB1 suppressed the genistein-induced increase in NAMPT expression and NAD^+^ levels in adipocytes. Genistein-bound PHB1 contributed to the stabilization of the transcription factor CCAAT/enhancer-binding protein β through activation of extracellular signal-regulated kinase, resulting in increased NAMPT expression at the transcriptional level. Genistein induced dephosphorylation of peroxisome proliferator-activated receptor at serine 273 and increased the insulin-sensitizing adipokine, adiponectin, in adipocytes, whereas the knockdown of NAMPT and PHB1 abolished these genistein-mediated effects. Our results proved the potential efficacy of nutraceuticals in promoting NAD^+^ levels and restoring metabolic function in adipocytes. Furthermore, we identified PHB1, localized to the plasma membrane, as a candidate target protein for increased expression of NAMPT in adipocytes. Overall, these findings will assist in developing NAD^+^ boosting strategies to alleviate the metabolic dysfunctions in adipose tissues.

**Significance Statement:** Increasing NAD^+^ levels is an important preventive strategy for maintaining metabolic function. Here, we showed that genistein, a nutraceutical, which increases NAD^+^ levels in adipocytes, increased NAD^+^ biosynthesis by upregulating nicotinamide phosphoribosyltransferase (NAMPT), a rate-limiting enzyme in the NAD^+^ biosynthesis pathway. Our findings also showed that genistein increased NAMPT expression by binding to prohibitin 1 in the plasma membrane. Genistein-induced increase in NAD^+^ levels promoted adiponectin expression, an insulin-sensitizing adipokine, in adipocytes. This study provides evidence that nutraceuticals, such as genistein, are effective in enhancing NAD^+^ biosynthesis in adipocytes and that PHB1 is a candidate target protein for increased expression of NAMPT to maintain metabolic functions in adipose tissues.

## Introduction

Obesity is a major global health challenge because of its association with systemic metabolic dysfunctions in different organs, leading to several diseases, including type 2 diabetes mellitus (T2D), cardiovascular diseases, and hypertension (1). Obesity-induced adipose tissue dysfunction leads to the dysregulated secretion of adipokines, such as adiponectin and adipsin, which are associated with insulin resistance (2).

Nicotinamide adenine dinucleotide (NAD^+^), a biological redox cofactor, plays vital roles in cellular energy metabolism and homeostasis in several organs. A decrease in NAD^+^ levels causes cellular dysfunction in adipocytes, hence, dysregulated secretion of adipokines, which is correlated with obesity and aging in rodents and humans (3-5). NAD^+^ is biosynthesized from its precursors, tryptophan, nicotinamide, and nicotinamide riboside. Nicotinamide phosphoribosyltransferase (NAMPT) is the rate-limiting enzyme in NAD^+^ biosynthesis and converts nicotinamide into nicotinamide mononucleotide (NMN), which is then rapidly converted to NAD^+^ by nicotinamide mononucleotide adenylyl transferase (6). Several studies have explored the potential of NMN treatment to increase NAD^+^ levels; however, its effects on insulin sensitivity are localized to skeletal muscles only (7,8). In contrast, the deletion of NAMPT in adipose tissue specifically decreases NAD^+^ levels in adipose tissue, resulting in insulin resistance (9,10). Furthermore, NAMPT inhibition suppresses adipocyte differentiation and decreases adiponectin secretion from adipocytes (11,12). These studies indicate that adipocyte NAD^+^ contributes to maintaining systemic metabolic functions; therefore, developing strategies to increase NAD^+^ levels (NAD^+^ boosting strategy) in adipocytes by inducing NAMPT expression could help alleviate metabolic dysfunctions.

Several studies have documented the pharmacological approaches to improve adipocyte function; however, their efficacy is not fully realized because of their side effects, such as weight gain and myocardial infarction (13). On the contrary, nutraceuticals, such as functional foods and dietary supplements, have been shown to alleviate adipose tissue dysfunctions without major side effects. Genistein (4′,5,7-trihydroxyisoflavone) is a nutraceutical derived from soybeans seeds, a major agricultural crop in the United States. Genistein has various physiological activities, and clinical studies have shown that genistein improves glucose metabolism and increases adiponectin levels in human trials (14,15). However, the molecular mechanisms of how genistein improves glucose metabolism and adiponectin levels in adipocytes remain unclear.

The metabolic function of adipocytes is mediated by several transcription factors, among which proliferator-activated receptor γ (PPARγ) and members of the CCAAT/enhancer-binding protein (C/EBP) family (C/EBPα and C/EBPβ) are the master regulators (16). PPARγ induces the expression of genes involved in glucose and lipid metabolism, such as adiponectin and adipsin, and inflammation (17). The phosphorylation of PPARγ, particularly at Ser273, is associated with obesity and insulin resistance (18,19). Furthermore, the overexpression of mutant PPARγ, which abolishes Ser273 phosphorylation, increases the expression of adiponectin (*ADIPOQ*) gene (18). In mice, it has been shown that some inhibitors of PPARγ phosphorylation at Ser237 increase insulin sensitivity (20,21), wherein increased NAD^+^ levels have been shown to suppress PPARγ phosphorylation at Ser273 (10). These findings suggest regulating PPARγ phosphorylation can alleviate insulin resistance and maintain metabolic functions in adipose tissues.

Based on the roles of genistein and NAD^+^ levels in increasing adiponectin levels and the maintenance of metabolic dysfunction, respectively, we hypothesized that the physiological activities of genistein are modulated via an increase in NAD^+^ levels in adipocytes. To test this hypothesis, here, we investigated the effect of genistein on NAD^+^ biosynthesis and explored the underlying mechanism. Our findings show that genistein increases NAD^+^ and adiponectin levels by inducing NAMPT. Furthermore, we uncovered that plasma membrane protein prohibitin 1 (PHB1) is involved in genistein-induced NAMPT expression. The findings provide insights into the role of genistein in maintaining adipocyte metabolic functions and can assist in developing nutraceutical-based approaches to overcome metabolic dysfunctions.

## Results

### Genistein promotes NAD^+^ biosynthesis by increasing NAMPT expression

We investigated whether genistein affects intracellular NAD^+^ levels and the expression of NAMPT in 3T3-L1 cells and adipose stromal cells (ASCs). Genistein treatment increased intracellular NAD^+^ (Fig. 1A) and intracellular NAMPT (iNAMPT) (Fig. 1B) levels in both adipocytes dose-dependently. Genistein-treated adipocytes showed a concentration-dependent increase in extracellular NAMPT (eNAMPT) levels in the culture supernatants (Fig. 1C), which contributes to NAD^+^ biosynthesis in other tissues. Genistein also increased the expression of *Nampt* at the transcriptional level (Fig. 1D). In contrast, genistein did not affect the expression of other NAD^+^ salvage pathway-related enzymes such as nicotinamide mononucleotide adenylyltransferase 1 (NMNAT)1, NMNAT2, and nicotinamide N-methyltransferase (NNMT) (Fig. S1). NAMPT knockdown reduced genistein-induced increases in NAD^+^ levels (Fig. 1E), suggesting that genistein increases NAD^+^ biosynthesis by increasing the expression of NAMPT.

**Figure 1.**
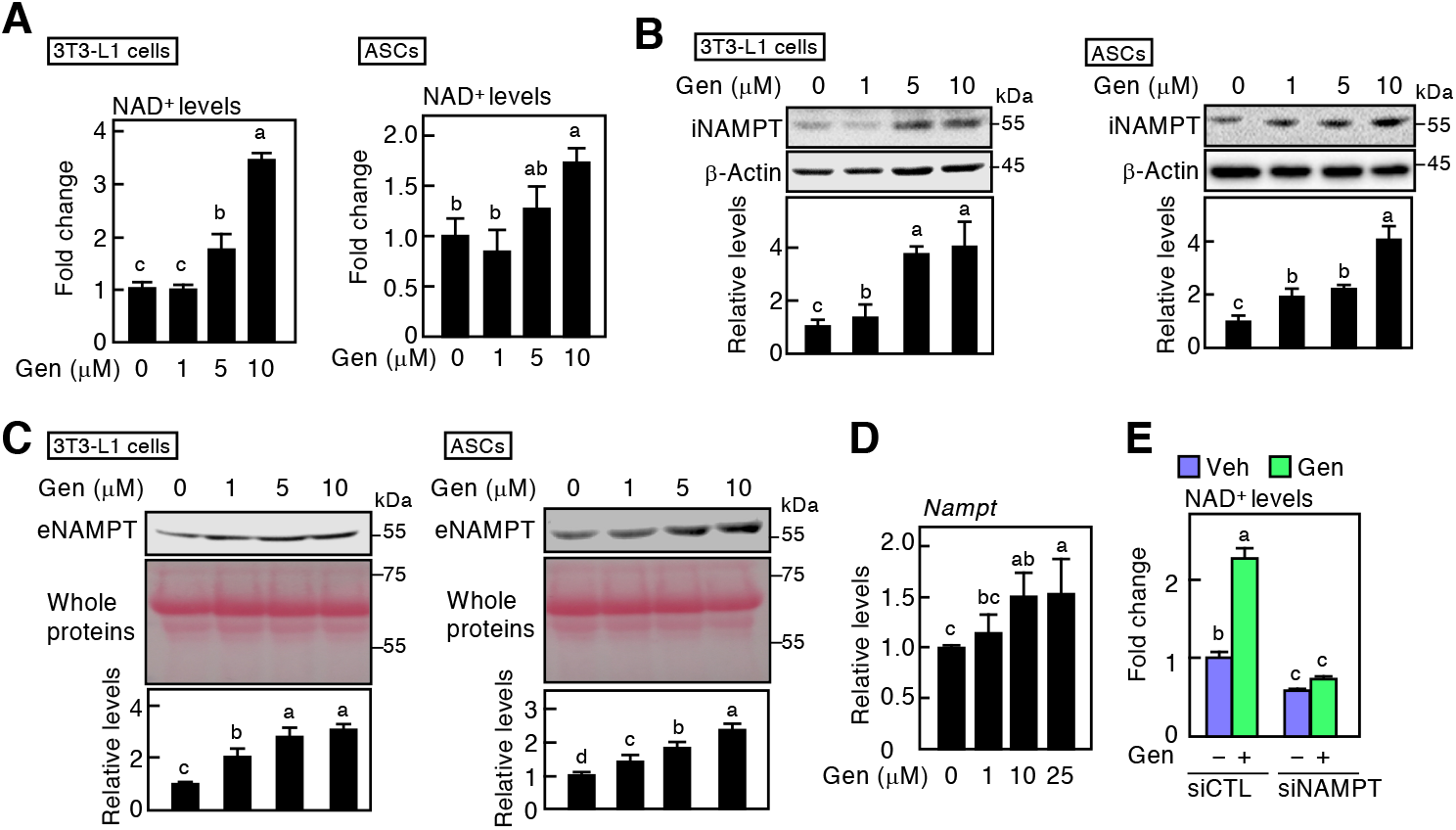
Effect of genistein on intracellular NAD^+^ and NAMPT expression levels. (A) Quantification of intracellular NAD^+^ levels in 3T3-L1 adipocytes after induction of differentiation with genistein (Gen) for seven days. (B) Western blotting of intracellular NAMPT (iNAMPT) in 3T3-L1 adipocytes (*left panels*) and ASCs (*right panels*) after induction of differentiation with Gen for seven days. The ratio of each band was normalized to that of β-actin. (C) Western blotting of extracellular NAMPT (eNAMPT) in 3T3-L1 adipocytes (*left panels*) and ASCs (*right panels*). Total protein was stained with Ponceau-S, and the ratio of each eNAMPT band was normalized to the total protein level. (D) *Nampt* gene expression in 3T3-L1 adipocytes treated with Gen for seven days. (E) Intracellular NAD^+^ levels in NAMPT knockdown 3T3-L1 adipocytes incubated in the presence (black bars) or absence (gray bars) of Gen (10 μM) for seven days. Data are representative of triplicate independent experiments presented as the mean ± SD (n = 3). Statistically significant differences (*p* < 0.05) are indicated using different letters— the difference is not statistically significant if two groups share at least one letter.

### PHB1 is involved in genistein-induced increase in NAMPT expression

To identify the genistein target proteins required for the induction of NAMPT expression, first, we determined the active site of the isoflavone involved in the induction of NAMPT expression by treating 3T3-L1 cells with genistein, 4′,7-dihydroxyisoflavone, or 5,7-dihydroxy-4′-methoxyisoflavone (4′-methylgenistein) (Fig. S2A). NAMPT levels were increased by genistein, and 4′,7-dihydroxyisoflavone, whereas 4′-methylgenistein did not affect NAMPT expression (Fig. S2B). Therefore, we used affinity beads to identify proteins that bind to genistein but not 4′-methylgenistein. Incubation of genistein-affinity beads with whole-cell extracts from 3T3-L1 adipocytes identified several bands of proteins specifically bound to genistein beads (Fig. 2A). Genistein-bound proteins were then digested into peptides and analyzed using quadrupole time-of-flight mass spectrometer. The predicted peptide sequences were searched in a mouse protein database in Uniprot, which revealed a sequence coverage of 43.4% (Fig. S2C the peptide sequences were characterized as PHB1(arrowhead bind in Fig. 2A). Subsequently, western blotting confirmed that PHB1 interacted with the genistein-affinity beads but not with the 4′-methylgenistein-affinity beads (Fig. 2B, *upper panels*). In contrast, the genistein beads did not bind to NAMPT (Fig. S2D), indicating that genistein did not directly affect the NAMPT protein via interaction with NAMPT. However, purified recombinant PHB1 interacted with genistein beads (Fig. 2B *bottom panels*).

**Figure 2.**
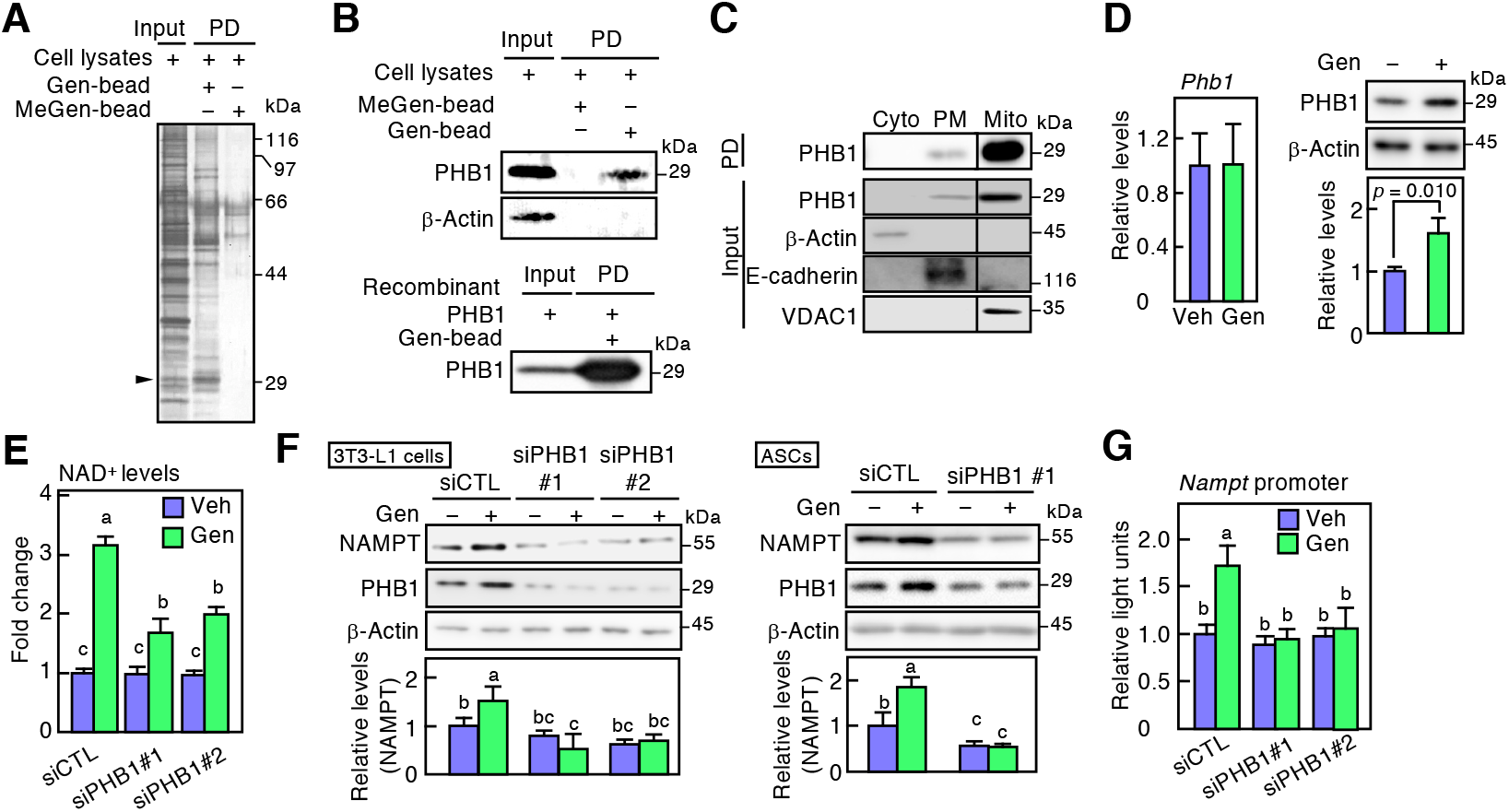
Identification of target protein for genistein-mediated regulation of NAMPT expression. (A) 3T3-L1 adipocyte lysates were incubated with the genistein-affinity beads (Gen-bead) or 4′-methylgenistein-affinity beads (MeGen-bead). Bead-bound proteins were collected by pull-down (PD) assay and subjected to SDS-PAGE, followed by silver staining. The arrow indicates the identified protein band. (B) Pull-down assay between Gen-bead and PHB1. 3T3-L1 adipocyte lysates (*top panels*) or recombinant His-tagged PHB1 (*bottom panels*) were incubated with Gen-bead or MeGen-bead. (C) Pull-down assay with Gen-bead and subcellular fractions in 3T3-L1 adipocytes. Proteins from the cytosol (Cyto), plasma membrane (PM), and mitochondria (Mito) were incubated with Gen-bead, followed by western blotting. (D) Analysis of *Phb1* expression levels in 3T3-L1 adipocytes treated with Gen (10 μM) for 24 h (*left panel*). Western blotting in 3T3-L1 adipocytes treated with Gen (10 μM) in the presence of cycloheximide (10 μg/mL) (*right panels*). The ratio of each band was normalized to that of β-actin. (E) Measurement of intracellular NAD^+^ levels in 3T3-L1 adipocytes treated with control siRNA (siCTL) or PHB1-specific siRNA (siPHB1). (F) Western blotting in PHB1 knockdown 3T3-L1 adipocytes (*left panels*) and ASCs (*right panels*). The ratio of each band was normalized to that of β-actin. (G) *Nampt* promoter activity in PHB1 knockdown 3T3-L1 adipocytes transiently transfected with pGL4-*Nampt-*Luc vector, followed by incubation with or without Gen in the presence of DMI. Data are representative of triplicate independent experiments presented as the mean ± SD (n = 3). Statistically significant differences (*p* < 0.05) are indicated using different letters—a common letter between the groups indicated that the difference is not statistically significant.

Next, we investigated the intracellular localization of the interaction between PHB1 and genistein. Incubation of subcellularly fractionated cell lysates from 3T3-L1 adipocytes with genistein beads that PHB1 was distributed in the mitochondria and plasma membrane. Furtehremore, genistein beads interacted with mitochondrial PHB1, as well as with plasma membrane-localized PHB1 (Fig. 2C). Subsequently, to examine the effect of genistein on PHB1, we analyzed the gene and protein expression of *PHB1* in genistein-treated adipocytes. Genistein did not affect the expression of *Phb1*; however, PHB1 expression increased in the presence of cycloheximide, a protein synthesis inhibitor (Fig. 2D), indicating that genistein upregulated the stability of PHB1. Evaluation of the effect of PHB1 on the genistein-induced increase in NAD^+^ levels revealed that PHB1 knockdown attenuated the genistein-induced increase in intercellular NAD^+^ (Fig. 2E) and NAMPT levels in adipocytes (Fig. 2F). Furthermore, we showed that genistein stimulated the *Nampt* promoter activity, which was suppressed by the knockdown of PHB1 (Fig. 2G). These results indicate that genistein upregulates the stability of PHB1 by binding directly to PHB1, resulting in increased expression of NAMPT at the transcriptional level.

### The interaction between genistein and PHB1 is important for genistein-induced NAMPT expression

PHB1 comprises an N-terminal transmembrane domain (NTD), SPFH domain (SPFHD), and C-terminal coiled-coil domain (CCD) (22,23). To identify the region of PHB1 that binds genistein, we constructed three PHB1 deletion mutants (Fig. 3A, *left panel*). Genistein beads were incubated with the recombinant PHB1 deletion mutants. The pull-down assay showed a physical interaction of genistein only with SPFHD (Fig. 3A, *right panel*). Furthermore, we showed that PHB1 lacking the NTD (PHB1ΔNTD) was localized in the cytoplasm rather than in the plasma membrane (Fig. 3B), indicating that the NTD of PHB1 is essential for its membrane localization (23). To determine whether plasma membrane-localized PHB1 was involved in the genistein-induced increase in NAMPT expression, we estimated the expression of NAMPT in 3T3-L1 adipocytes stably expressing PHB1ΔNTD. Although genistein increased PHB1ΔNTD levels, NAMPT levels were not affected by genistein in PHB1ΔNTD-expressing adipocytes (Fig. 3C), indicating that NTD is not involved in genistein-induced NAMPT expression. Therefore, to identify the region of PHB1 that binds genistein, genistein beads were incubated with two deletion mutants of recombinant SPFHD in PHB1. We found that genistein bound to amino acids 55–94 of PHB1 (Fig. 3D).

**Figure 3.**
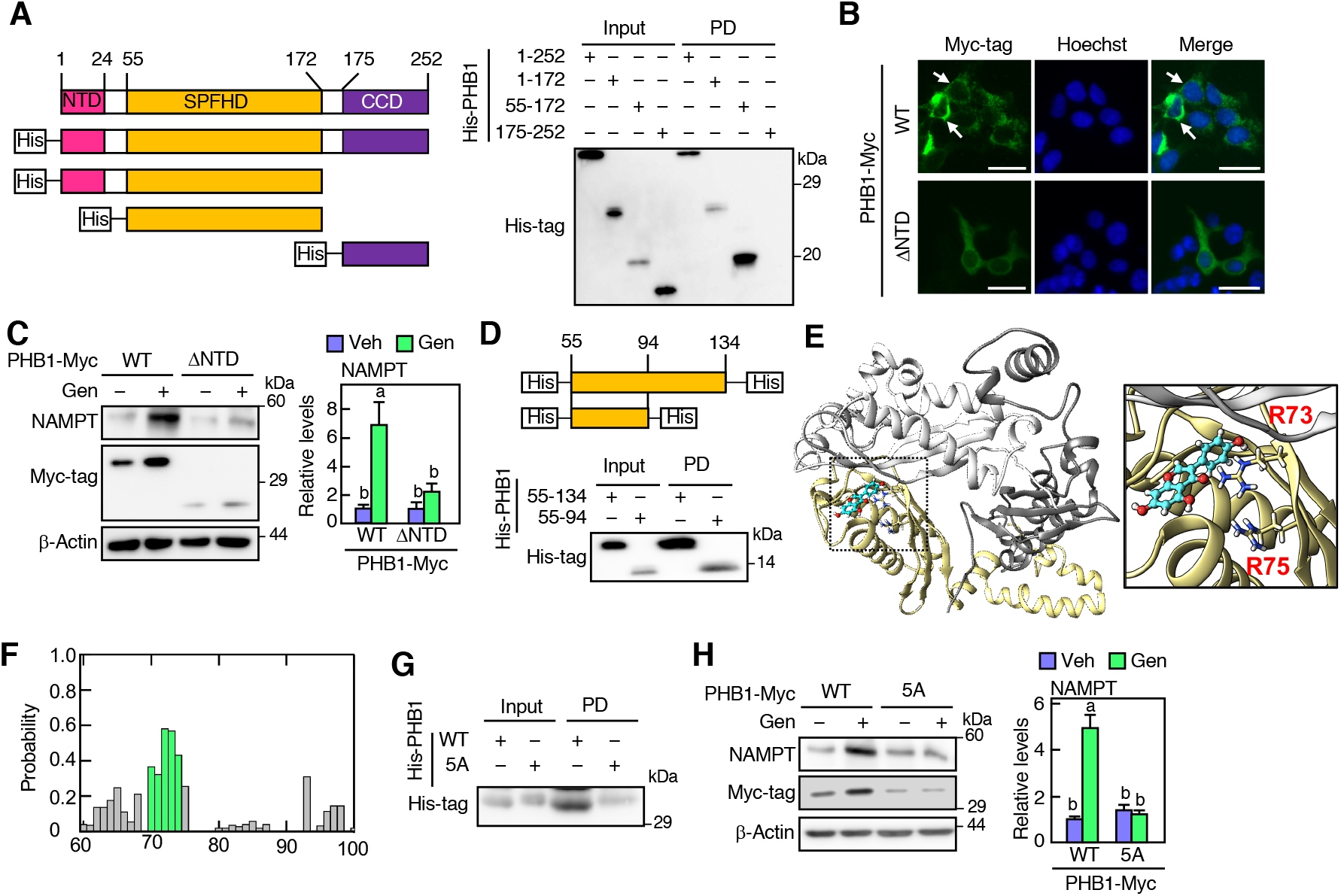
Determination of interacting region between genistein and PHB1. (A) Schematic representation of recombinant PHB1 fragments (*left panels*); NTD, N-terminal transmembrane domain; SPFHD, SPFH domain; CCD, C-terminal coiled-coil domain. Pull-down (PD) assay of recombinant His-PHB1 and genistein-affinity beads (Gen-bead). Pulled-down proteins were analyzed using western blotting (*right panels*). (B) Immunofluorescence staining of PHB1-Myc (green) in 3T3-L1 cells. Nuclei were stained with Hoechst 33258 (blue); Scale bar, 25 μm. (C) Western blotting in 3T3-L1 adipocytes expressing mutant PHB1. Cells were infected with Myc-tag infused PHB1(wild-type; WT) or PHB1(ΔNTD) expression lentivirus, followed by induction of differentiation in the presence or absence of Gen (10 μM) for 7 days. The ratio of each NAMPT band was normalized to that of β-actin. (D) Schematic representation of the PHB1 fragment in the SPFH domain (*upper panels*). Pull-down assay with Gen-bead and recombinant His-PHB1 fragments (*lower panel*). (E) Best docking pose of Gen with PHB1 trimer (yellow, white, and gray complex). The Gen molecule is represented as a ball-stick model with cyan carbon atoms. The first (yellow), second (white), and third (gray) chains of PHB1 trimer are represented as ribbons. In the first chain, the side chains of R73 and R75 were displayed as stick models. (F) Interaction sites of Gen docking to PHB1 target region (60–94 amino acid residues) by ensemble docking simulation. The x-axis denotes the residue number of PHB1. The green bars indicate amino acid residues from 70 to 74 in the PHB1 protein. (G) Pull-down assay with Gen-bead and recombinant His-tag infused PHB1(WT) or mutant PHB1. (H) Protein expression in 3T3-L1 adipocytes expressing mutant PHB1. Cells were infected with Myc-tag-infused PHB1(WT) or PHB1(5A) expression lentivirus, followed by induction of differentiation in the presence or absence of Gen for seven days. The ratio of each NAMPT band was normalized to that of β-actin. Data are representative of triplicate independent experiments presented as the mean ± SD (n = 3). Statistically significant differences (*p* < 0.05) are indicated using different letters—a common letter between the groups indicated that the difference is not statistically significant.

Subsequently, we performed computational molecular dynamics and docking to develop structural models of PHB1 using an ensemble docking simulation to investigate the structural basis of the interaction between genistein and PHB1. Docking simulations showed that PHB1 formed a trimer at amino acid residues 55–94. The best position of genistein on the PHB1 trimer suggested that the positively charged residues R73 and R75 of PHB1 interacted with the oxygen atoms of genistein, which could have contributed to stabilizing the complex (Fig. 3E). The probability of a docking site was higher near positively charged residues (70–74 amino acid residues) (Fig. 3F). To determine whether the five amino acid residues of PHB1 at positions 70– 74 are required for the interaction with genistein, we constructed an expression vector of mutant *Phb1* in which the five amino acids of PHB1 (at 70–74) were replaced with Ala [PHB1(5A)]. The pull-down assay showed that PHB1(5A) interacted less with genistein than wild-type (WT) PHB1 (Fig. 3F). Furthermore, genistein did not affect NAMPT levels in PHB1(5A)-expressing adipocytes (Fig. 3G). These results indicate that genistein strongly interacts with PHB1 at amino acid residues 70–74, and this interaction is important for genistein-enhanced NAMPT expression.

### Genistein enhances the C/EBPβ-mediated promoter activity of *Nampt*

Our previous study showed that cAMP-induced C/EBPβ stimulates *Nampt* promoter activity (11). Therefore, we hypothesized that cAMP and C/EBPβ are involved in genistein-stimulated *Nampt* promoter activity. To test this hypothesis, we examined promoter activity in 3T3-L1 cells.

Genistein alone did not affect *Nampt* promoter activity. In contrast, it enhanced the activity in the presence of isobutylmethylxanthine (IBMX, an inhibitor of cAMP degradation) (Fig. 4A).

**Figure 4.**
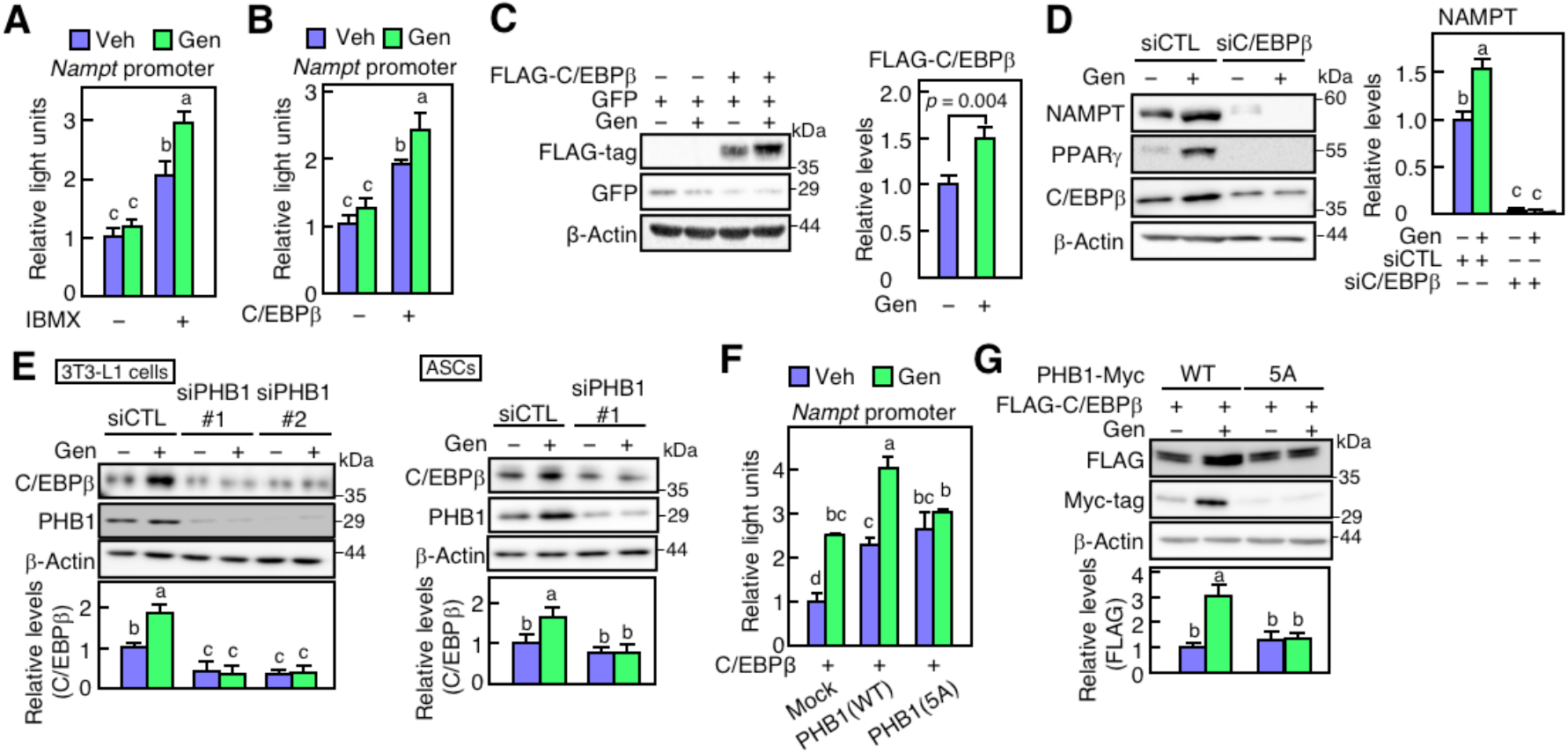
Effect of genistein and PHB1 on C/EBPβ-stimulated promoter activity of *Nampt* gene. (A) *Nampt* promoter activity in 3T3-L1 adipocytes. After transfection, the cells were incubated with genistein (Gen; 10 μM) in the presence or absence of IBMX. (B) *Nampt* promoter activity in 3T3-L1 adipocytes transfected with Myc-C/EBPβ expression vector. (C) Protein expression of exogenous C/EBPβ in 3T3-L1 adipocytes. Cells were transiently transfected with FLAG-C/EBPβ or GFP (control) expression vectors. The ratio of each FLAG-C/EBPβ band was normalized to that of β-actin. (D) Western blotting in C/EBPβ knockdown 3T3-L1 adipocytes. Cells were transfected with siRNA for control (siCTL) or C/EBPβ specific (siC/EBPβ). The ratio of each NAMPT band was normalized to that of β-actin. (E) Protein expression in PHB1 siRNA (siPHB1) treated 3T3-L1 adipocytes (*left panels*) or ASCs (*right panels*). The ratio of each C/EBPβ band was normalized to that of β-actin. (F) *Nampt* promoter activity in 3T3-L1 adipocytes transfected C/EBPβ and *PHB1* expression vectors. (G) Protein expression of exogenous C/EBPβ in 3T3-L1 adipocytes expressing mutant *PHB1*. Cells were infected with Myc-tag-infused *PHB1*(wild-type; WT) or *PHB1*(5A) expression lentivirus, followed by transient transfection with FLAG-C/EBPβ expression vectors. The ratio of each FLAG-C/EBPβ band was normalized to that of β-actin. Data are representative of triplicate independent experiments presented as the mean ± SD (n = 3). Statistically significant differences (*p* < 0.05) are indicated using different letters—a common letter between the groups indicated that the difference is not statistically significant.

Furthermore, in C/EBPβ-expressing 3T3-L1 cells, genistein also stimulated its promoter activity (Fig. 4B), indicating that genistein enhances C/EBPβ-mediated *Nampt* promoter activity. Further investigation of the effects of genistein on C/EBPβ levels showed that increased levels of exogenous C/EBPβ but not those of exogenous GFP (Fig. 4C). Moreover, genistein increased endogenous C/EBPβ levels and knockdown of C/EBPβ blunted the effect of genistein on the induction of the NAMPT expression (Fig. 4D). However, genistein did not affect the mRNA levels of *Cebpb* (Fig. S3A). Next, we investigated whether genistein modulated C/EBPβ levels by interacting with PHB1—the knockdown of PHB1 suppressed genistein-induced C/EBPβ levels in 3T3-L1 adipocytes and ASCs (Fig. 4E). As shown in Fig. 3E, PHB1(5A) binds genistein less tightly than WT PHB1. Moreover, WT PHB1 stimulated C/EBPβ-mediated *Nampt* promoter activity, and genistein further enhanced *Nampt* promoter activity (Fig. 4F). In contrast, genistein did not enhance *Nampt* promoter activity in PHB1(5A)- and PHB1ΔNTD-expressing conditions (Figs. 4F and S3B). Exogenous C/EBPβ levels were increased by genistein in WT PHB1-expressing conditions but not in PHB1(5A)-expressing conditions. Furthermore, the expression of PHB1(5A) was lower than that of WT PHB1, and genistein only increased the expression of WT PHB1 (Fig. 4G), suggesting that genistein regulates the stability of C/EBPβ by binding to PHB1. Overall, our results indicate that genistein enhances NAMPT expression by increasing the stability of C/EBPβ and interacting with PHB1.

### Genistein stimulates ERK signaling-induced increase in stability of C/EBPβ by binding to PHB1

Next, we investigated the mechanism of how genistein stimulates C/EBPβ stability via PHB1. We showed that genistein induced the phosphorylation of ERK, and the knockdown of PHB1 blunted it in 3T3-L1 adipocytes and ASCs (Figs. 5A and S4A). In contrast, the ERK inhibitor PD98059 suppressed genistein-increased C/EBPβ expression (Figs. 5B and S4B), *Nampt* promoter activity (Fig. 5C), and intercellular NAD^+^ levels (Fig. 5D) in the presence of genistein. Furthermore, the stability of C/EBPβ is regulated by its phosphorylation at threonine 188 (Thr188) (24,25).

**Figure 5.**
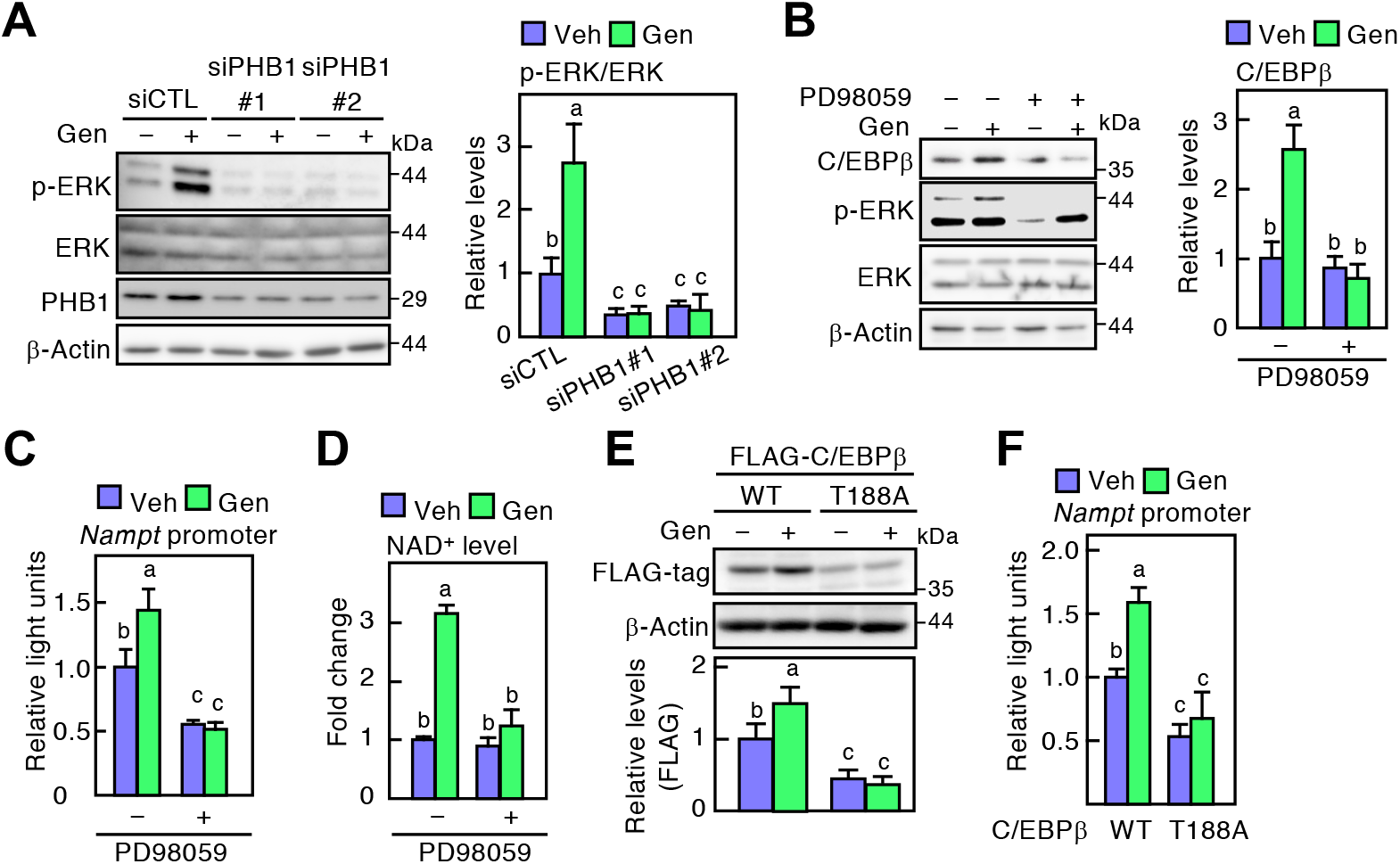
Association of genistein-increased C/EBPβ protein levels and ERK signaling. (A) Phosphorylation levels of ERK in PHB1-specific siRNA (siPHB1)-treated 3T3-L1 adipocytes treated with genistein (Gen; 10 μM). (B) Western blotting in 3T3-L1 adipocytes treated with Gen and ERK inhibitor PD98059 (20 μM). The ratio of each C/EBPβ band was normalized to that of β-actin. (C) *Nampt* promoter activity in 3T3-L1 adipocytes treated with ERK inhibitor. (D) Measurement of intracellular NAD^+^ levels in 3T3-L1 adipocytes treated with Gen and PD98059. (E) Western blotting in exogenous C/EBPβ-expressed 3T3-L1 cells. Cells were transiently transfected with FLAG-C/EBPβ(wild-type; WT) or FLAG-C/EBPβ(T188A) expression vectors. The ratio of each FLAG-C/EBPβ band was normalized to that of β-actin. (F) *Nampt* promoter activity in exogenous C/EBPβ mutant-expressed 3T3-L1 adipocytes. Data are representative of triplicate independent experiments presented as the mean ± SD (n = 3). Statistically significant differences (*p* < 0.05) are indicated using different letters—a common letter between the groups indicated that the difference is not statistically significant.

Therefore, to assess the relationship between the phosphorylation at Thr188 and the stability of C/EBPβ, we generated a C/EBPβ mutant with Thr188 replaced by Ala (T188A). The level of C/EBPβ (T188A) was lower than that of WT C/EBPβ in the absence of genistein, suggesting that Thr188 is important for C/EBPβ stability. Furthermore, WT C/EBPβ level was increased by genistein, whereas that of C/EBPβ (T188A) did not change by genistein (Fig. 5E). Furthermore, the C/EBPβ (T188A)-mediated *Nampt* promoter activity was lower than that mediated by WT C/EBPβ wherein genistein had no effect on *Nampt* promoter activity in C/EBPβ (T188A)-expressing cells (Fig. 5F). Taken together, our results demonstrated that genistein increases C/EBPβ levels through its phosphorylation via the PHB1-ERK signaling pathway.

### Acetylation levels of PPARγ are regulated by genistein-enhanced NAMPT expression

Next, we investigated the effect of the genistein-induced increase in NAD^+^ levels via NAMPT expression in adipocytes. NAD^+^ is a substrate for the SIRT-mediated deacetylation reaction, and NAMPT knockout causes a reduction in adiponectin levels through an increase in the lysine acetylation of PPARγ (10). Therefore, it is suggested that NAMPT-increased NAD^+^ induces adiponectin expression through the increase in PPARγ deacetylation. Immunoprecipitation experiments showed that genistein decreased acetylated PPARγ levels in adipocytes (Fig. 6A). Furthermore, knockdown of *Nampt* and *Phb1* suppressed genistein-mediated deacetylation of PPARγ (Fig. 6B). To evaluate the effect of genistein-induced deacetylation of PPARγ on adiponectin expression, we constructed a mutant form of PPARγ (Lys268Gln and Lys293Gln; 2KQ) that is not deacetylated by SIRT1 (26). The results showed that genistein increased the adiponectin levels in WT PPARγ-but not in PPARγ(2KQ)-expressing adipocytes (Fig. 6C), indicating that genistein promotes the deacetylation by inducing NAMPT expression.

**Figure 6.**
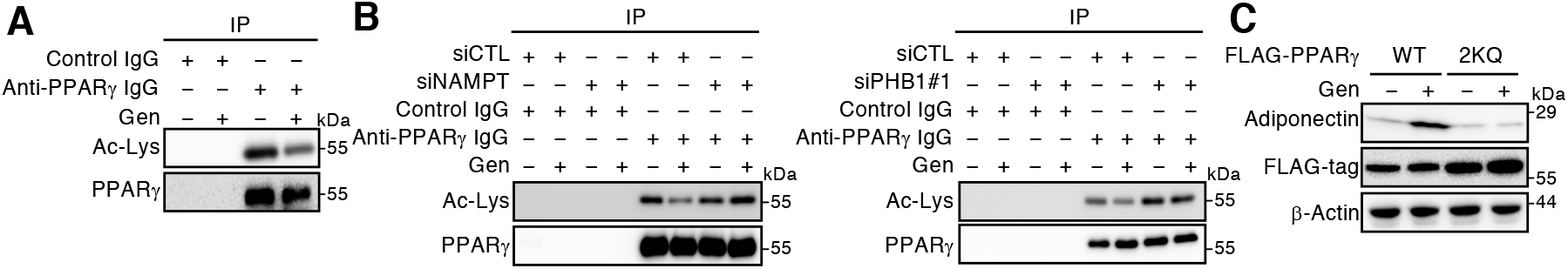
Involvement of NAMPT in genistein-induced deacetylation of PPARγ. (A) Analysis of acetylated PPARγ in 3T3-L1 adipocytes treated with genistein (Gen; 10 μM), and PPARγ was immunoprecipitated (IP) using anti-PPARγ IgG. (B) Acetylation of PPARγ in NAMPT-(*left panels*) or PHB1-knockdown (*right panels*) 3T3-L1 adipocytes treated with Gen, and PPARγ was immunoprecipitated using anti-PPARγ IgG. (C) Analysis of PPARγ phosphorylation in acetylation mimics PPARγ-expressed 3T3-L1 adipocytes. Cells were infected with FLAG-PPARγ(wild-type; WT) or FLAG-PPARγ(K268Q and K293Q; 2KQ) lentivirus and differentiated. FLAG-PPARγ was immunoprecipitated. *D*, Western blotting of adiponectin in PPARγ(2KQ)-expressed 3T3-L1 cells. The cells were infected with FLAG-PPARγ(WT) or FLAG-PPARγ(S273E; SE) lentivirus. Data representative of triplicate independent experiments.

### Genistein induces adiponectin expression by decreasing the expression of phosphorylated PPARγ

The assessment of the effects of genistein on the phosphorylation of PPARγ revealed that genistein decreased Ser273 phosphorylation in WT PPARγ but not in PPARγ(2KQ) (Fig. 7A). Furthermore, genistein decreased the Ser273 phosphorylation of endogenous PPARγ in a dose-dependent manner (Fig. 7B). In contrast, protein levels of adiponectin and total PPARγ were increased by genistein. Concordantly, genistein increased the expression of *Adipoq* and *Adn* (encoding adipsin), whereas the expression of other PPARγ target genes, such as *Fabp4* and *Cd36* (encoding fatty acid binding protein 4 and cluster of differentiation 36, respectively), was not affected by genistein (Fig. 7C). We hypothesized that genistein affects the phosphorylation by increasing the expression of NAMPT and NAD^+^. To test this hypothesis, we analyzed the phosphorylation of PPARγ at Ser273 in *Nampt*-knockdown adipocytes. NAMPT knockdown abolished the genistein-induced reduction in phosphorylated PPARγ levels (Figs. 7D and S5A). In addition, *NAMPT* knockdown decreased genistein-induced increases in *Adipoq* and *Adn* expression (Fig. 7E). On the contrary, FK866, an enzymatic inhibitor of NAMPT, suppressed the genistein-induced increase in adiponectin protein and gene expression (Fig. S5B and C). To determine whether genistein-mediated dephosphorylation of PPARγ is involved in the induction of adiponectin expression, 3T3-L1 adipocytes were treated with WT PPARγ or PPARγ(Ser273Glu; SE), which mimics phosphorylated PPARγ at Ser273. Genistein increased adiponectin protein levels in WT PPARγ-expressed cells, but it did not affect those in PPARγ(SE)-expressing cells (Fig. 7F). Subsequently, we determined whether PHB1 is involved in genistein-mediated dephosphorylation of PPARγ. PHB1 knockdown restored the genistein-induced decrease in the phosphorylated PPARγ, whereas the ERK inhibitor suppressed the genistein-induced dephosphorylation of PPARγ (Fig. 7G). Furthermore, genistein-induced adiponectin expression was suppressed by PHB1 knockdown and ERK inhibitor treatment (Fig. 7H). These results indicate that genistein-induced NAMPT increased adiponectin expression through the induction of PPARγ dephosphorylation.

**Figure 7.**
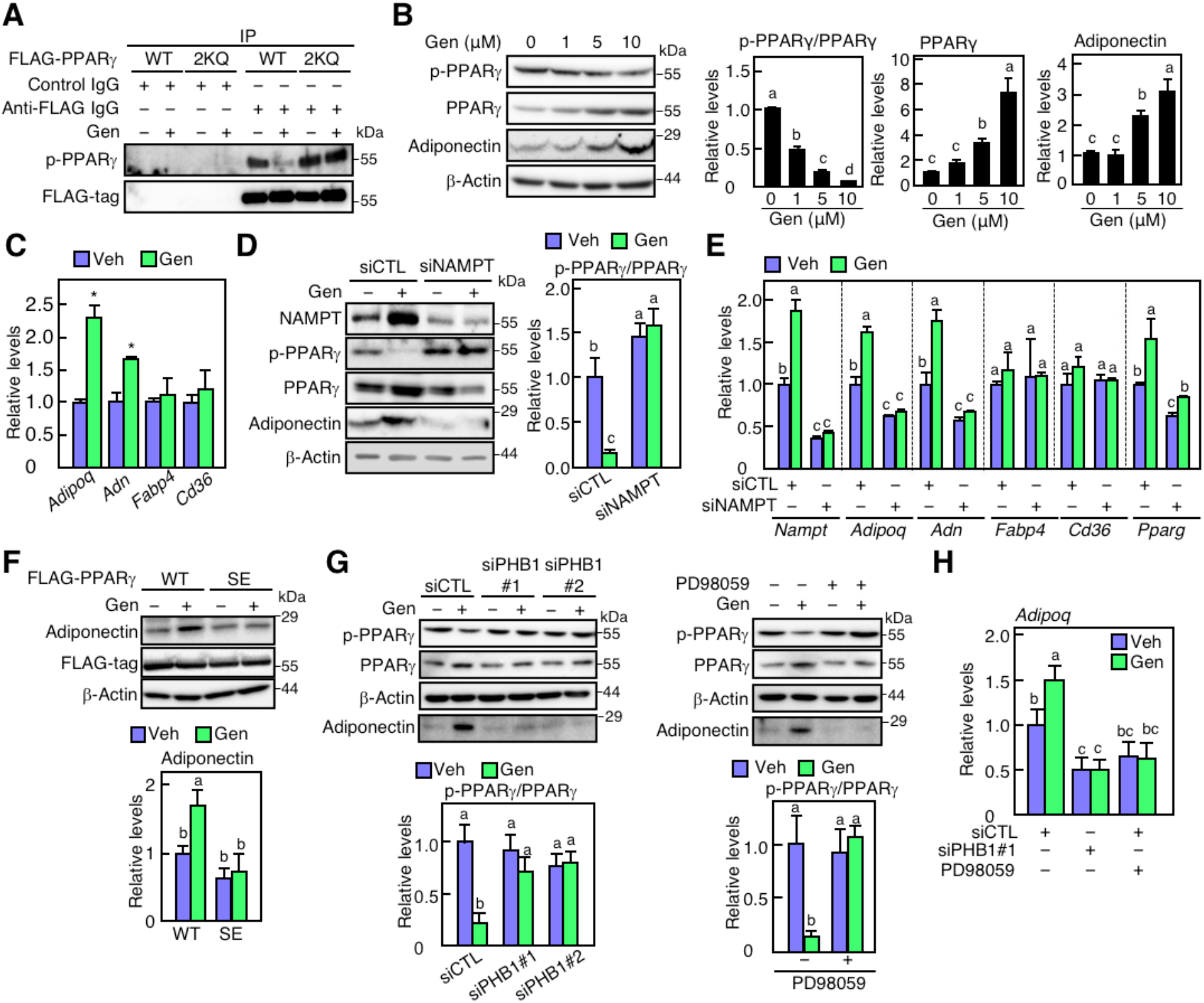
Effect of genistein on phosphorylation of PPARγ through the upregulation of NAMPT. (A) Phosphorylation levels of PPARγ (wild-type; WT) PPARγ (K268Q and K293Q; 2KQ) in 3T3-L1 adipocytes. Cells were differentiated in the presence of genistein (Gen; 10 μM) for 7 days. (B) Protein expression in 3T3-L1 adipocytes treated with Gen. The ratio of each band was normalized to that of β-actin. (C) qPCR analysis of PPARγ target genes in 3T3-L1 treated with Gen. (D) Western blotting in NAMPT knockdown 3T3-L1 adipocytes. Cells were transfected with siRNA for control (siCTL) or NAMPT specific (siNAMPT). (E) qPCR analysis in NAMPT-knocked down 3T3-L1 adipocytes. (F) Western blotting of adiponectin in phosphorylation mimics PPARγ-expressed 3T3-L1 cells. Cells were infected with FLAG-PPARγ(WT) or FLAG-PPARγ(S273E; SE) lentivirus. The ratio of each adiponectin band was normalized to that of β-actin. (G) Protein expression in 3T3-L1 adipocytes treated with PHB1 siRNA (siPHB1). The ratio of each p-PPARγ band was normalized to that of PPARγ. (H) Western blotting in 3T3-L1 adipocytes treated with Gen and ERK inhibitor PD98059 (20 μM). The ratio of each p-PPARγ band was normalized to that of PPARγ. Data are representative of triplicate independent experiments presented as the mean ± SD (n = 3). In (*C*), statistically, significant differences are indicated by asterisks (\**p* < 0.05 vs. Veh). In others, statistically significant differences (*p* < 0.05) are indicated using different letters—a common letter between the groups indicated that the difference is not statistically significant.

### Genistein administration increases NAMPT expression and NAD^+^ biosynthesis *in vivo*

To verify whether the effects of genistein obtained *in vitro* could be observed *in vivo*, we orally administered genistein to mice for 14 days and measured the amount of NAD^+^ in the adipose tissues. We observed a significant increase in NAD^+^ levels in the perigonadal adipose tissue, which was classified as the visceral adipose tissue (Fig. 8A). Analysis of the expression levels of each protein showed that genistein treatment increased the expression of NAMPT and adiponectin in perigonadal adipose tissues (Fig. 8B), suggesting that NAMPT is involved in the genistein-induced increase in NAD^+^ levels. In contrast, genistein did not increase NAMPT levels in subcutaneous and interscapular adipose tissues or skeletal muscle (Fig. S6A). Genistein treatment decreased phosphorylated PPARγ and increased adiponectin levels (Fig. 8B); however, it did not affect body weight or fat mass (Fig. S6B). These results strongly suggest that genistein supplementation enhances NAD^+^ biosynthesis *in vivo*.

**Figure 8.**
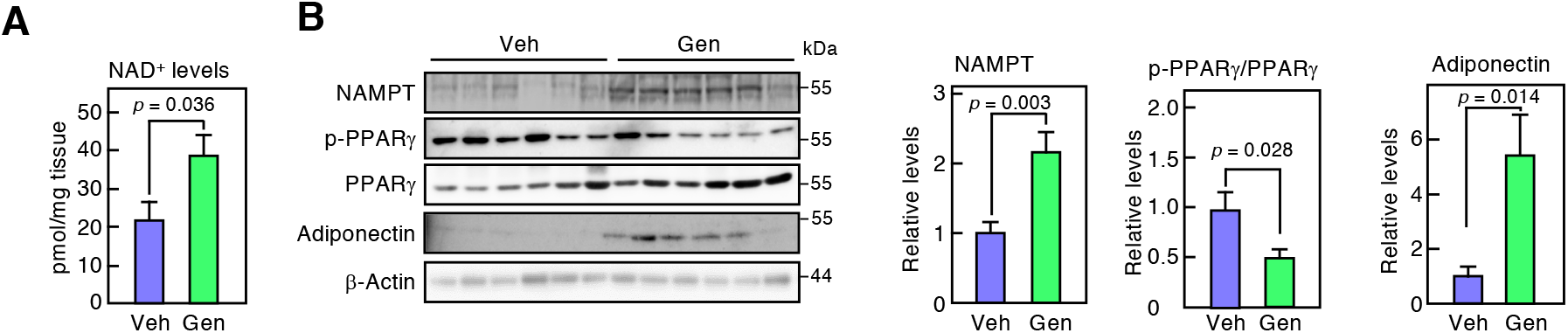
Effect of genistein administration on NAD^+^ levels and NAMPT expression in adipose tissues. (A) Quantification of NAD^+^ levels in visceral adipose tissues of mice after 14 days of oral administration of vehicle (Veh) or genistein (Gen). (B) Western blotting in mouse visceral adipose tissue (*left panels*). The ratio of each band was normalized to that of β-actin (*right panels*). Error bars represent the mean ± SEM (n = 6), and statistically significant differences are indicated using asterisks (\**p* < 0.05 vs. Veh).

## Discussion

This study provides evidence that nutraceutical strategies can improve the NAD^+^ levels in adipocytes and elucidate its underlying mechanism. Our results showed that genistein treatment (1) increased NAD^+^ biosynthesis by upregulating NAMPT expression, (2) enhanced the stability of C/EBPβ via PHB1 and ERK signaling, and (3) promoted adiponectin expression by inducing deacetylation of PPARγ. These results provide important insights into the maintenance of adipose tissue metabolism by increasing NAD^+^ levels using nutraceuticals.

Several studies have demonstrated the health-promoting effects of genistein, both *in vivo* and *in vitro*; however, its target proteins remain unknown. As genistein is structurally similar to estrogen, it has been speculated to bind to estrogen (ERs) and nuclear receptors and induce transcriptional activation (27,28). Studies have also shown that genistein inhibits breast cancer growth and obesity in ovariectomized rats via ER signaling (29,30). Nevertheless, some studies have shown that genistein is ER-independent (31–33). For instance, in skeletal muscle cells, the expression of the β2-adrenergic receptor is upregulated by genistein, which is independent of ER (32). Genistein increases adiponectin expression by inhibiting tyrosine kinases in synovial fibroblasts (33) and suppresses ATP synthesis by binding to the mitochondrial protein ANT2 in preadipocytes (31). In addition, molecular docking studies have shown that genistein binds to c-Jun-NH2-terminal kinase 1 (JNK1) and suppresses TNF-α-mediated downregulation of adiponectin by inhibiting the JNK1 in mature adipocytes (34). Here, we demonstrated that genistein binds directly to PHB1 in adipocytes. These studies indicate that genistein exerts multiple biological effects by targeting various proteins. Our results identified a genistein target protein that may explain its functional properties in adipocytes.

Knockdown of PHB1 reduced the genistein-enhanced NAD^+^ biosynthesis, and PHB1ΔNTD attenuated the genistein-induced increased expression of NAMPT. *Phb* encodes two protein isoforms, PHB1 and PHB2, mainly localized in the inner mitochondrial membrane, nucleus, cytoplasm, and plasma membrane (35). In the inner mitochondrial membrane, PHB1 and PHB2 form a heterooligomeric complex that stabilizes the mitochondrial network (35), and PHBs expression is negatively correlated with mitochondrial oxidative stress levels (36). In the plasma membrane, PHB1 acts as a transmembrane receptor that activates downstream signaling. At the T cell surface, PHB1 activates the Rac1 and Raf-ERK signaling cascades by acting as an adapter that recognizes viral peptides, thereby inhibiting viral entry and triggering inflammatory responses (37). Moreover, the small compound PDD005 binds to PHB1 in the plasma membrane and reduces neuroinflammation by decreasing IL-1β in aged mice (38). Chronic treatment with PDD005 increased PHB1 levels in the brains of aged mice, suggesting that PDD005 contributes to PHB1 stability. We found that genistein-activated ERK signaling was abolished by PHB1 knockdown, suggesting that genistein increases *Nampt* expression by binding to PHB1 on the plasma membrane than on the inner mitochondrial membrane in adipocytes. Although PHB1 is highly expressed in the endothelial cell membranes of human and mouse adipose tissue (39), its function in adipocytes remains elusive. It has been reported that PHB1 knockdown causes mitochondrial fragmentation and decreases the activity of oxidative phosphorylation in 3T3-L1 cells (40). However, similar results were obtained with PHB1 and PHB2 knockdown. These results indicated that PHBs localized in the mitochondrial inner membrane could be involved in mitochondrial maintenance in adipocytes. Our previous study showed that genistein binds to mitochondrial proteins (31), suggesting that genistein translocates to the mitochondria and may affect the function of mitochondrial PHB1. However, this should be verified in future studies. These results indicate that plasma membrane-localized PHB1 is essential for the genistein-induced increase in NAMPT expression in adipocytes and demonstrate the function of PHB1 in regulating gene expression by activating intracellular signaling in adipocytes.

The genistein-mediated *Nampt* promoter activity was suppressed by an ERK inhibitor. Earlier, we have shown that NAMPT is a target gene of C/EBPβ in adipocytes (11). C/EBPβ is phosphorylated at Thr188 by the MAPK/ERK signaling pathway and CDK2 during the early stage of adipogenesis. Using phosphorylation at Thr188 as a marker, C/EBPβ is phosphorylated at Ser184 and Thr179 by GSK3β, which induces a conformational change in C/EBPβ resulting in increased DNA-binding activity (41). Post-translational modification regulates C/EBPβ through DNA-binding activity and protein stability during adipogenesis (24,42,43). In addition, the stability of C/EBPβ(T188A) is lower than that of WT C/EBPβ (25,44). Consistent with these findings, we showed that genistein increased expression of the WT C/EBPβ but not C/EBPβ(T188A). Together with the data showing that plasma membrane PHB1 activates the ERK signaling pathway, these results indicate that ERK-mediated stabilization of C/EBPβ is critical for genistein-increased *Nampt* expression.

Genistein induces PPARγ dephosphorylation by enhancing NAD^+^ biosynthesis, while full agonists of PPARγ prevent and ameliorate insulin resistance and T2D; they also cause secondary effects, including weight gain, congestive heart failure, and reduced bone mineral density (45,46). This is because PPARγ upregulates the expression of genes involved in lipogenesis and fatty acid synthesis. It has been shown that the p-PPARγ(Ser273) levels are increased in the adipose tissue of mice fed a high-fat diet and insulin-resistant mice (18,19).

Furthermore, mouse models with a double-allele mutation in S273A exhibited a more insulin-sensitive phenotype than WT mice fed a high-fat diet (47). It has also been shown that PPARγ(S273A) overexpression increased the expression of *Adipoq, Adn*, and *Lep*, but not those of other target genes, such as *Fabp4, Fasn, Cd36* in PPARγ(S273A)-expressing cells (18), suggesting that PPARγ regulates the expression of its target genes through phosphorylation at Ser273. Therefore, the focus should be on the development of deactivating compounds that regulate post-translational modifications of PPARγ. The non-agonist PPARγ ligands SR1664 and UHC1 inhibit the CKD5-mediated phosphorylation of PPARγ at Ser293 (48,49). These ligands do not affect PPARγ-mediated adipogenesis but increase the expression of *Adipoq* and *Adn*, resulting in improved insulin sensitivity in high-fat diet-fed mice. On the contrary, a ligand-independent regulatory mechanism of PPARγ phosphorylation has been proposed. Lys268 and Lys293 of PPARγ are targets for deacetylation by SIRT1. Qiang *et al*. (26) suggested that the hyperacetylation of Lys268 and Lys293of PPARγ triggers phosphorylation at Ser273. Concordant with these studies, we found that genistein-induced reduction of p-PPARγ(Ser273) levels disappeared in acetylated mimetic PPARγ at Lys268 and Lys293. Moreover, the deletion of NAMPT increases p-PPARγ(Ser273) levels and decreases *Adipoq* expression (10). In contrast, NMN administration restored NAMPT-suppressed expression, indicating that enhancement of NAD^+^ biosynthesis indirectly regulates the dephosphorylation of PPARγ via its deacetylation. These results demonstrate that genistein indirectly regulates post-translational modifications of PPARγ by enhancing NAD^+^ biosynthesis rather than acting as a ligand for PPARγ. Taken together, we speculated that genistein could alleviate insulin resistance and T2D by regulating PPARγ dephosphorylation without secondary effects such as weight gain.

Our study demonstrated the effects of genistein on NAMPT and adiponectin expression *in vitro* and *in vivo*. Although other researchers have shown that NAD^+^ elevation suppresses body weight and fat mass gain (8,50), we did not observe a reduction after 14 days of genistein administration, possibly due to the short duration of genistein administration. Furthermore, when fed a chow diet for 12 weeks, there was no difference in body weight or fat mass gain between WT and adipose tissue-specific *Nampt*-knockout mice (51). Therefore, further studies using *Nampt*-knockout mice fed a high-fat diet are needed to establish the health-promoting effects of genistein supplementation.

In conclusion, this study provides insights into enhancing NAD^+^ biosynthesis in adipocytes by genistein, a nutraceutical. Although future studies are required to address the effect of genistein-mediated activation of NAD^+^ biosynthesis in adipose tissue on systemic glucose and lipid metabolism in obese or aged animal models, it is important to note that nutraceuticals are effective in enhancing NAD^+^ biosynthesis in adipocytes and that PHB1, which is localized to the plasma membrane, is a candidate target protein for increased expression of NAMPT.

## Materials and methods

### Animal experiments

Eight-week-old male ICR mice were purchased from Japan SLC, Inc. (Shizuoka, Japan) and housed under controlled temperature (20 ± 3°C) with a 12 h light/dark cycle and free access to food and water. After one week of acclimatization, the mice were randomly divided into genistein (LC laboratories) and vehicle groups (n = 6 per group) and administered with genistein suspended in corn oil (50 mg/kg body weight) or vehicle (corn oil), respectively by oral gavage once daily for 14 days. At the end of the experiment, mice were sacrificed under isoflurane inhalation anesthesia. Adipose tissues were collected, and samples were stored at –80°C until analysis. All animal experiment protocols were approved by the Institutional Animal Care and Use Committee of Shinshu University Animal Experimentation Regulations (Approval Nos. 300046 and 019024). The study has been designed and performed in accordance with the ARRIVE guidelines (52).

### Cell culture

Methods for culture and induction of differentiation of murine 3T3-L1 cells (No. IFO50416, JCRB Cell Bank, Osaka, Japan) and immortalized mouse ASCs are described previously (31). The cells were differentiated into adipocytes for 7 days. Genistein was added to the medium from day 2 of the initiation of differentiation.

### siRNA

Target sequences of the siRNA duplexes are listed in Table S1. All siRNA duplexes were synthesized by Sigma-Aldrich. The control siRNA (siCTL) was purchased from Sigma-Aldrich (MISSION siRNA Universal Negative Control#1). The duplexes (20 nM) were transfected into cells using the Lipofectamine RNAiMAX reagent (Invitrogen, Carlsbad, CA, USA) and Opti-MEM (Thermo Fisher Scientific, Lafayette, CO, USA) for 24 h according to the manufacturer’s protocol.

### Intracellular NAD^+^ assay

The intracellular NAD^+^ in 3T3-L1 cells or ASCs was measured using the NAD^+^/NADH-Glo assay kit (Promega) following the manufacturer’s instructions. Adipose tissue NAD^+^ levels were analyzed using an enzymatic cycling assay (51). Briefly, adipose tissue (100 mg) was homogenized in 400 μL perchloric acid (0.6 M). The supernatant was diluted 10-fold in 100 mM Na_2_HPO_4_ (pH 8.0), and the sample was reacted with 2% ethanol, 90 U/mL alcohol dehydrogenase, 130 mU/mL diaphorase, 10 μM resazurin, 10 μM flavin mononucleotide, and 10 mM nicotinamide for 4 h at 37°C. Resorufin accumulation was measured using fluorescence excitation at 530 nm and emission at 590 nm.

### Identification of genistein-binding proteins

The preparation of isoflavone-fixed beads and the pull-down assay of genistein-bound proteins were previously described (31). Bead-bound proteins were subjected to SDS-PAGE, followed by silver staining. Selected gel bands that did not bind to the 4’-methylgenistein beads were subjected to in-gel digestion with trypsin (MS grade; Promega). Peptide fragments were analyzed using a Q-TOF/MS, and the processed peptide data were searched against the mouse protein database in UniProt (www.uniprot.org).

### Pull-down assay

The pET-30a-PHB1 vector encoding His-tagged mouse PHB1 was transformed into *Escherichia coli* strain BL21 Star (DE3; Stratagene). The resulting N-terminal His-tagged recombinant PHB1 was purified using Ni-Sepharose 6 Fast Flow (GE Healthcare). Recombinant His-tagged PHB1 was incubated with genistein beads (0.5 mg) in pull-down buffer for 4 h at 4°C, and the beads were then washed five times with wash buffer. Bead-bound proteins were subjected to SDS-PAGE and analyzed by western blotting using an anti-His-tag antibody.

### Luciferase reporter assay

3T3-L1 cells were transiently transfected with reporter vectors (pGL4-*Nampt*-Luc (11), and pRL-SV40; control reporter vector; Promega) or Myc-C/EBPβ expression vector (pLVSIN-Myc-C/EBPβ) using Lipofectamine 3000 (Thermo Fisher Scientific) for 24 h. After replacing the medium with fresh medium, the cells were incubated with IBMX and genistein (10 μM) for 24 h. Firefly and Renilla luciferase activities were measured using a dual-luciferase reporter assay kit and a GloMax 20/20 Luminometer (Promega). Transfection efficiency was normalized to that of Renilla luciferase. Data are expressed as relative light units.

### Co-immunoprecipitation

The co-immunoprecipitation assay was performed as described previously (53). Briefly, 3T3-L1 cells were differentiated into adipocytes with or without genistein (10 μM) for 7 days. Cells were lysed in denaturing cell extraction buffer, and the cell lysates were incubated with mouse monoclonal anti-PPARγ IgG, anti-FLAG IgG, or control IgG at 4°C overnight, followed by incubation with protein G-Sepharose resin (50% slurry; GE Healthcare, Waukesha, WI) at 4°C for 2 h. The resin was washed three times with lysis buffer, and the proteins bound to the resin were separated by SDS-PAGE and analyzed using western blotting.

### Statistical analyses

Data are expressed as mean ± SD or ± SEM. Significant differences between the two groups were determined using Student’s *t*-test. Data from more than three groups were analyzed using one- or two-way analysis of variance with Tukey’s post-hoc test. All statistical analyses were performed using the JMP statistical software version 11.2.0 (SAS Institute, Cary, NC, USA). Statistical significance was set at *p* < 0.05.

Other materials and methods are described in the Supplementary Infromation (SI).

## Acknowledgments

This work was supported by JSPS KAKENHI (grant number: 20K05922 and 23K05109) for scientific research (to T.M.), the Mishima Kaiun Memorial Foundation (http://www.mishima-kaiun.or.jp/), and the Shinshu Foundation for Promotion of Agricultural and Forest Science to T.M. Protein detection, identification, and qPCR analyses were conducted at the Research Center for Support to Advanced Science at Shinshu University. We would like to thank Editage (www.editage.jp) for English language editing.

## Author Contributions

S.W. and T.M. conceptualized the study. S.W. and T.M. designed the study. S.W., T.M., R.H., K.U., and I.T. performed the investigations. S.W., T.M., and K.U. analyzed the data; S.W. and T.M. wrote the original draft; S.W., T.M., and U.K. wrote, reviewed, and edited the paper; S.N. and S.K. contributed to resources.

## References

1 O’Neill S, O’Driscoll L. 2015. Metabolic syndrome: a closer look at the growing epidemic and its associated pathologies. Obes Rev. 16:1–12.

2 Klöting N, et al. 2010. Insulin-sensitive obesity. Am J Physiol Endocrinol Metab. 299:E506–E515.

3 Yoshino J, Mills KF, Yoon MJ, Imai S. 2011. Nicotinamide mononucleotide, a key NAD(+) intermediate, treats the pathophysiology of diet- and age-induced diabetes in mice. Cell Metab. 14:528–536.

4 Yoshino J, Baur JA, Imai S. 2018. NAD+ intermediates: the biology and therapeutic potential of NMN and NR. Cell Metab. 27:513–528

5 Cantó C, Menzies KJ, Auwerx J. 2015. NAD(+) metabolism and the control of energy homeostasis: a balancing act between mitochondria and the nucleus. Cell Metab. 22:31–53.

6 Xie N, et al. 2020. NAD+ metabolism: pathophysiologic mechanisms and therapeutic potential. Signal Transduct Target Ther. 5:227.

7 Yoon MJ, et al. 2015. SIRT1-mediated eNAMPT secretion from adipose tissue regulates hypothalamic NAD+ and function in mice. Cell Metab. 21:706–717.

8 Stromsdorfer KL, et al. 2016. NAMPT-mediated NAD(+) biosynthesis in adipocytes regulates adipose tissue function and multi-organ insulin sensitivity in mice. Cell Rep. 16:1851–1860.

9 Mitani T, et al. 2020. Intracellular cAMP contents regulate NAMPT expression via induction of C/EBPβ in adipocytes. Biochem Biophys Res Commun. 522:770–775.

10 Okabe K, et al. 2020. NAD+ metabolism regulates preadipocyte differentiation by enhancing α-ketoglutarate-mediated histone H3K9 demethylation at the PPARγ promoter. Front Cell Dev Biol. 8:586179.

11 Yoshino M, et al. 2021. Nicotinamide mononucleotide increases muscle insulin sensitivity in prediabetic women. Science. 372:1224–1229.

12 Mills KF, et al. 2016. Long-term administration of nicotinamide mononucleotide mitigates age-associated physiological decline in mice. Cell Metab. 24:795–806.

13 Nissen S and Wolski K. 2007. Effect of rosiglitazone on the risk of myocardial infarction and death from cardiovascular causes. N Engl J Med. 356:2457–2471.

14 Atteritano M, et al. 2007. Effects of the phytoestrogen genistein on some predictors of cardiovascular risk in osteopenic, postmenopausal women: a two-year randomized, double-blind, placebo-controlled study. J Clin Endocrinol Metab. 92:3068–3075.

15 Squadrito F, et al. 2013. Genistein in the metabolic syndrome: results of a randomized clinical trial. J Clin Endocrinol Metab. 98:3366–3374.

16 Lefterova MI, et al. 2008. PPARγ and C/EBP factors orchestrate adipocyte biology via adjacent binding on a genome-wide scale. Genes Dev. 22:2941–2952.

17 Festuccia WT, et al. 2009. Depot-specific effects of the PPARγ agonist rosiglitazone on adipose tissue glucose uptake and metabolism. J Lipid Res. 50:1185–1194.

18 Choi JH, et al. 2010. Anti-diabetic drugs inhibit obesity-linked phosphorylation of PPARγ by Cdk5. Nature. 466:451–456.

19 Banks AS, et al. 2015. An ERK/Cdk5 axis controls the diabetogenic actions of PPARγ. Nature. 517:391–395.

20 Amato AA, et al. 2012. GQ-16, a novel peroxisome proliferator-activated receptor γ (PPARγ) ligand, promotes insulin sensitization without weight gain. J Biol Chem. 287: 28169–28179.

21 Choi SS, et al. 2016. PPARγ antagonist Gleevec improves insulin sensitivity and promotes the browning of white adipose tissue. Diabetes. 65:829–839.

22 Tatsuta T, Model K, Langer T. 2005. Formation of membrane-bound ring complexes by prohibitins in mitochondria. Mol Biol Cell. 16:248–259.

23 Winter A, Kämäräinen O, Hofmann A. 2007. Molecular modeling of prohibitin domains. Proteins. 68:353–362.

24 Tang QQ, et al. 2005. Sequential phosphorylation of CCAAT enhancer-binding protein β by MAPK and glycogen synthase kinase 3β is required for adipogenesis. Proc Natl Acad Sci U S A. 102:9766–9771.

25 Zhang Y, et al. 2012. Phosphorylation prevents C/EBPβ from the calpain-dependent degradation. Biochem Biophys Res Commun. 419:550–555.

26 Qiang L, et al. 2012. Brown remodeling of white adipose tissue by SirT1-dependent deacetylation of Pparγ. Cell. 150:620–632.

27 Adlercreut CH, et al. 1995. Soybean phytoestrogen intake and cancer risk. J Nutr. 125:757S–770S.

28 Ogawa M, et al. 2017. Daidzein down-regulates ubiquitin-specific protease 19 expression through estrogen receptor β and increases skeletal muscle mass in young female mice. J Nutr Biochem. 49:63–70.

29 Hsieh CY, Santell RC, Haslam SZ, Helferich WG. 1998. Estrogenic effects of genistein on the growth of estrogen receptor-positive human breast cancer (MCF-7) cells in vitro and in vivo. Cancer Res. 58:3833–3838.

30 Shen HH, et al. 2019. Genistein ameliorated obesity accompanied with adipose tissue browning and attenuation of hepatic lipogenesis in ovariectomized rats with high-fat diet. J Nutr Biochem. 67:111–122.

31 Ikeda T, Watanabe S, Mitani T. 2022. Genistein regulates adipogenesis by blocking the function of adenine nucleotide translocase-2 in the mitochondria. Biosci Biotechnol Biochem. 86:260–272.

32 Chikazawa M, Sato R. 2018. Identification of a novel function of resveratrol and genistein as a regulator of β2-adrenergic receptor expression in skeletal muscle cells and characterization of promoter elements required for promoter activation. Mol Nutr Food Res. 62:e1800530.

33 Relic B, et al. 2009. Genistein induces adipogenesis but inhibits leptin induction in human synovial fibroblasts. Lab Invest. 89:811–822.

34 Yanagisawa M, et al. 2012. Genistein and daidzein, typical soy isoflavones, inhibit TNF-α-mediated downregulation of adiponectin expression via different mechanisms in 3T3-L1 adipocytes. Mol Nutr Food Res. 56:1783–1793.

35 Mishra S, Murphy LC, Nyomba BLG, Murphy LJ. 2005. Prohibitin: a potential target for new therapeutics. Trends Mol Med. 11:192–197.

36 Chai RR, et al. 2017. Prohibitin involvement in the generation of mitochondrial superoxide at complex I in human sperm. J Cell Mol Med. 21:121–129.

37 Chowdhury I, Thompson WT, Thomas K. 2014. Prohibitins role in cellular survival through Ras-Raf-MEK-ERK pathway. J Cell Physiol. 229:998–1004.

38 Guyot AC, et al. 2020. A small compound targeting prohibitin with potential interest for cognitive deficit rescue in aging mice and tau pathology treatment. Sci Rep. 10:1143.

39 Kolonin MG, Saha PK, Chan L, Pasqualini R, Arap W. 2004. Reversal of obesity by targeted ablation of adipose tissue. Nat Med. 10. 625–632.

40 Liu D, et al. 2012. Mitochondrial dysfunction and adipogenic reduction by prohibitin silencing in 3T3-L1 cells. PLoS One. 7:e34315.

41 Guo L, Li X, Tang QQ. 2015. Transcriptional regulation of adipocyte differentiation: a central role for CCAAT/enhancer-binding protein (C/EBP) β. J Biol Chem. 290:755–761.

42 Km JW, Tang QQ, Li X, Lane MD. 2007. Effect of phosphorylation and S-S bond-induced dimerization on DNA binding and transcriptional activation by C/EBPβ. Proc Natl Acad Sci U S A. 104:1800–1804.

43 Li X, et al. 2007. Role of cdk2 in the sequential phosphorylation/activation of C/EBPβ during adipocyte differentiation. Proc Natl Acad Sci U S A. 104. 11597–11602.

44 Takai T, et al. 2018. Casein kinase 2 phosphorylates and stabilizes C/EBPβ in pancreatic β cells. Biochem Biophys Res Commun. 497:451–456.

45 Nesto RW, et al. 2004. Thiazolidinedione use, fluid retention, and congestive heart failure: a consensus statement from the American Heart Association and American Diabetes Association. Diabetes Care. 27:256–263.

46 Ali AA, et al. 2005. Rosiglitazone causes bone loss in mice by suppressing osteoblast differentiation and bone formation. Endocrinology. 146:1226–1235.

47 Hall JA, et al. 2020. Obesity-linked PPARγ S273 phosphorylation promotes insulin resistance through growth differentiation factor 3. Cell Metab. 32:665–675.

48 Choi JH, et al. 2011. Antidiabetic actions of a non-agonist PPARγ ligand blocking Cdk5-mediated phosphorylation. Nature. 477:477–481.

49 Chio SS, et al. 2014. A novel non-agonist peroxisome proliferator-activated receptor γ (PPARγ) ligand UHC1 blocks PPARγ phosphorylation by cyclin-dependent kinase 5 (CDK5) and improves insulin sensitivity. J Biol Chem. 289:26618–26629.

50 Cantó C, et al. 2012. The NAD(+) precursor nicotinamide riboside enhances oxidative metabolism and protects against high-fat diet-induced obesity. Cell Metab. 15:838–847.

51 Nielsen KN, et al. 2018. NAMPT-mediated NAD+ biosynthesis is indispensable for adipose tissue plasticity and development of obesity. Mol Metab. 11:178–188.

52 de Sert NP, et al. 2020. The ARRIVE guidelines 2.0: Updated guidelines for reporting animal research. PLOS Biol. 18:e3000410

53 Mitani T, et al. 2017. Theobromine suppresses adipogenesis through enhancement of CCAAT-enhancer-binding protein β degradation by adenosine receptor A1. Biochim Biophys Acta Mol Cell Res. 1864:2438–2448.

