## Supplementary materials and methods for "Genistein enhances NAD^+^ biosynthesis by binding to Prohibitin 1 and upregulating nicotinamide phosphoribosyltransferase in adipocytes"

**This PDF file includes:**

Supplementary text
Figures S1 to S6
Tables S1 to S2
SI References 5

**Appendix: Materials and methods**

**Plasmid and lentiviruses.** Three tandem FLAG and Myc tags were inserted into the pLVSIN-CMV Pur vector (Takara Bio, Shiga, Japan) and termed pLVSIN-FLAG and pLVSIN-Myc, respectively. Mouse *Pparg* cDNA was amplified using PCR; *Pparg* cDNA encoding a *Pparg* mutant with Lys to Gln substitutions at 268 and 293 positions and a Ser to Glu substitution at position 273 was generated. These *Pparg* cDNAs were inserted into pLVSIN-FLAG to construct the mutant *Pparg* expression vectors: pLVSIN-FLAG-PPARγ wild-type (WT), pLVSIN-FLAG-PPARγ (2KQ; K268Q and K293Q), and pLVSIN-FLAG-PPARγ (SE; S273E), respectively. Mouse prohibitin 1 (*Phb1*) cDNA was amplified using PCR, and *Phb1* cDNA encoding a *Phb1* mutant with Arg-Ser-Arg-Pro-Arg for Ala substitution at positions 70 to 74 amino acids and deletion of the N-terminal transmembrane domain (NTD) was generated. These *Phb1* cDNAs were inserted into pLVSIN-Myc to construct the mutant *Phb1* expression vectors: pLVSIN-Myc-PHB1 (WT), pLVSIN-Myc-PHB1(5A), and pLVSIN-Myc-PHB1ΔNTD, respectively. The *Phb1* cDNA was subcloned into the pET-30a vector, yielding a *Phb1* expression vector with six tandem N-terminal His-tags (pET-30a-PHB1). Deletion mutants of *PHB1* (1–172, 55–172, 175–252, 55–134, and 55–94 amino acids) were amplified using PCR and subcloned into the pET-30a vector. For lentivirus production, HEK293 FT cells (Takara Bio, Ohtsu, Japan) were transfected with *PHB1*- or *Pparg* -expressing vectors and a high titer lentiviral packaging mix (Takara Bio, Shiga, Japan) using TransIT-293 reagent (Takara Bio) following the manufacturer's instructions. After 48 h, the viral supernatant was collected and filtered. The 3T3-L1 cells were incubated overnight with the viral supernatant supplemented with 8 µg/mL polybrene (Sigma Aldrich).

**Western blotting.** The collected 3T3-L1 cells and ASCs were lysed in lysis buffer (50 mM Tris-HCl, pH 7.4, containing 150 mM NaCl, 0.5% Nonidet-P40, 10 mM sodium pyrophosphate, 2 mM EDTA, and a protease inhibitor cocktail [Nacalai Tesque, Kyoto, Japan]). The cell lysates were subjected to sodium dodecyl sulfate-polyacrylamide gel electrophoresis and analyzed by western blotting using the following primary antibodies: mouse monoclonal antibodies [anti-PPARγ (clone; E-8; Santa Cruz Biotechnology, Santa Cruz, CA) and anti-β-actin (clone; C4; Santa Cruz Biotechnology), anti-acetylated Lysine (clone; 15G10; BioLegend, San Diego, CA), anti-FLAG (clone; M2; Sigma-Aldrich), anti-Myc (clone; MC045; Nacalai Tesque, and anti-His-tag (clone; GHis; MBL Co., Ltd., Nagoya, Japan)]; rabbit polyclonal antibodies [anti-C/EBPβ (Bethyl, Montgomery, TX), anti-adiponectin and p-PPARγ (Ser273) (Bioss antibody Inc., Woburn, MA), anti-ERK1/2 and anti-p-ERK1/2 (Thr202/Tyr204) (Cell Signaling Technology, Beverly, CA), and anti-prohibitin 1 and anti-NAMPT (GeneTex, San Antonio, TX)]. After incubation with primary antibodies for 2 h at 4°C, the blots were incubated with horseradish peroxidase-conjugated secondary antibodies for 2 h at 4°C, followed by Immunostar LD (Wako, Osaka, Japan). Chemiluminescence was detected using LAS500 (GE Healthcare).

**Docking genistein on a conformational ensemble of PHB1.** The all-atom molecular dynamics simulation was conducted for a trimer model of PHB1 (60-233 residue) predicted using the SWISS-MODEL server (1). We used the ff14SB force field for PHB1 and TIP3P for the water molecules. The explicit solvent contained 100 mM and neutralized the system. After energy minimization and equilibration, the production run was simulated for 500 ns at 310 K. Calculations were performed using the Amber2016 and AmberTools packages (2). Snapshots were recorded every 10 ns; 50 different conformations were obtained and used for ensemble docking simulations. The molecular geometries of genistein were optimized with B3LYP/6-31G(d) using the Gaussian 09 suite (3). The restrained electrostatic potential charges on the genistein atoms were determined using AmberTools antechamber. The genistein model was docked to the first chain of the PHB1 trimer using MyPresto (ver.5.0) Sievgene program (4). Docking simulations were performed for 50 PHB1 conformations, and the best scores and positions for each conformation were saved. The best positions were then analyzed to calculate the interaction sites between genistein and PHB1.

**Quantitative real-time PCR (qPCR).** cDNAs were synthesized using RevaTra Ace (Toyobo, Ohtsu, Japan) and subjected to qPCR using the primers listed in Table S2. qPCR was performed using the KAPA SYBR Green Master Mix (NIPPON Genetics Co., Ltd., Tokyo, Japan) and a two-step PCR method on a Thermal Cycler Dice real-time system (Takara Bio). Relative mRNA levels for each gene were calculated using the $2^{-\Delta\Delta Ct}$ method, and data were normalized to that of *Rn18S* (18S rRNA) as an endogenous control.

**Immunofluorescence microscopy.** Immunofluorescence analysis was performed as previously described (5). Briefly, 3T3-L1 cells were transfected with the Myc-PHB1 expression vector. The cells were fixed, permeabilized, and incubated with rabbit anti-c-Myc, followed by Alexa Fluor 488-conjugated secondary anti-rabbit IgG. The nuclei were stained with Hoechst 332581 and observed under a fluorescence microscope (FV1000-D; Olympus Optical Co. Ltd., Tokyo, Japan).

**SI References**


1. Waterhouse A, et al. 2018. SWISS-MODEL: homology modelling of protein structures and complexes. *Nucleic Acids Res.* 46:W296-303.
2. Case DA. Betz RM. Botello-Smith W, Cerutti DS, Cheatham TE III, Darden TA, Duke RE, Giese TJ, Gohlke H, Goetz AW, Homeyer N, Izadi S, Janowski P, Kaus J, Kovalenko A, Lee TS, LeGrand S, Li P, Lin C, Luchko T, Luo R, Madej B, Mermelstein D, Merz KM, Monard G, Nguyen H, Nguyen HT, Omelyan I, Onufriev A, Roe DR, Roitberg A, Sagui C, Simmerling CL, Swails J, Walker RC, Wang J, Wolf RM, Wu X, Xiao L, York DM, Kollman PA. AMBER 2016, University of California: San Francisco, 2016.
3. Frisch, M.J.; Trucks, G.W.; Schlegel, H.B.; Scuseria, G.E.; Robb, M.A.; Cheeseman, J.R.; Scalmani, G.; Barone, V.; Petersson, G.A.; Nakatsuji, H.; et al. Gaussian 09 Revision D. 01; Gaussian Inc.: Wallingford, CT, USA, 2009.
4. Fukunishi Y, Mikami Y, Nakamura H. 2005. Similarities among receptor pockets and among compounds: analysis and application to in silico ligand screening. *J Mol Graph Model.* 24:34-45.
5. Tanaka E, et al. 2022. Theobromine enhances the conversion of white adipocytes into beige adipocytes in a PPARγ activation-dependent manner. *J Nutr Biochem.* 100:108898.
