## Supplementary Table 1 for "Genistein enhances NAD^+^ biosynthesis by binding to Prohibitin 1 and upregulating nicotinamide phosphoribosyltransferase in adipocytes"

**Table S1.** Table showing the list of target sequences for siRNA.

| Target name | Accession Number | Sequences (5´-3´) |
|---|---|---|
| C/EBPβ | NM_009883 | GAGCGACGAGTACAAGAT |
| NAMPT | NM_021524 | GCGCTTTGCTACAGAAGTTAA |
| PHB1#1 | NM_118993 | GCTGTCATCTTTGACCGAT |
| PHB1#2 | NM_118993 | GCTTCCTCGTATCTACACC |
