## Supplementary Table 2 for "Genistein enhances NAD^+^ biosynthesis by binding to Prohibitin 1 and upregulating nicotinamide phosphoribosyltransferase in adipocytes"

**Table S2.** Table showing the list of primer sequences for qPCR.

| Gene | Accession Number | Forward primer (5´-3´) | Reverse primer (5´-3´) |
|---|---|---|---|
| *Rn18S* | NR_003278 | GTAACCCGTTGAACCCCATT | CCATCCAATCGGTAGTAGCG |
| *Adipoq* | NM_009605 | GAACTTGTGCAGGTTGGATG | TGCATCTCCTTTCTCTCCCT |
| *Adn* | NM_013459 | CATGCTCGTCATCCGTCAC | CACAGAGTCGTCATCCGTCAC |
| *Cd36* | NM_001159558 | ATGGGCTGTGATCGGAACTG | TTTGCCACGTCATCTGGGTTT |
| *Cebpb* | NM_009883 | GGGGTTGTTGATGTTTTTGG | CGAAACGGAAAAGGTTCTCA |
| *Fabp4* | NM_024406.3 | AAGGTGAAGAGCATCATAACCT | TCACGCCTTTCATAACACATTCC |
| *Nampt* | NM_021524 | CGGTTCTGGTGGCGCTTTGCT | AAGTTCCCCGCTGGTGTCCTA |
| *Nmnat1* | NM_001356357 | GTGCCCAACTTGTGGAAGAT | CAGCACATCGGACTCGTAGA |
| *Nmnat2* | NM_175460 | CCGTCTCATCATGTGTCAGC | ACACACTGCAGGTTGTCTGC |
| *Nnmt* | NM_001311062 | GGAATCTGGCTTCACCTCCAA | GGAATCTGGCTTCACCTCCAA |
| *Pparg* | NM_001127330 | AGAAGGAACACTTGTCAGCG | TCAGCCACTTGAGTGTCCTC |
| *Phb1* | NM_118993 | TCCCTTGGGTACGAAACCAATTA | TGTGATATTGACGTTCTGCAAGTCT |
