## Supplementary Figure 1 for "Genistein enhances NAD^+^ biosynthesis by binding to Prohibitin 1 and upregulating nicotinamide phosphoribosyltransferase in adipocytes"

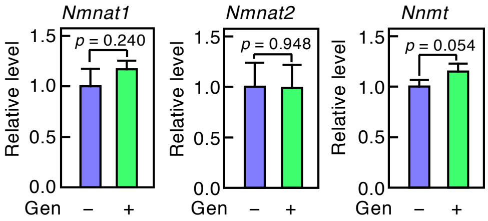


**Fig. S1.** qPCR analysis of $NAD^+$ biosynthesis associated genes in 3T3-L1 adipocytes after induction of differentiation with genistein (Gen; 10 μM) for 7 days. Data are representative of triplicate independent experiments presented as the mean ± SD (n = 3).
