## Supplementary Figure 2 for "Genistein enhances NAD^+^ biosynthesis by binding to Prohibitin 1 and upregulating nicotinamide phosphoribosyltransferase in adipocytes"

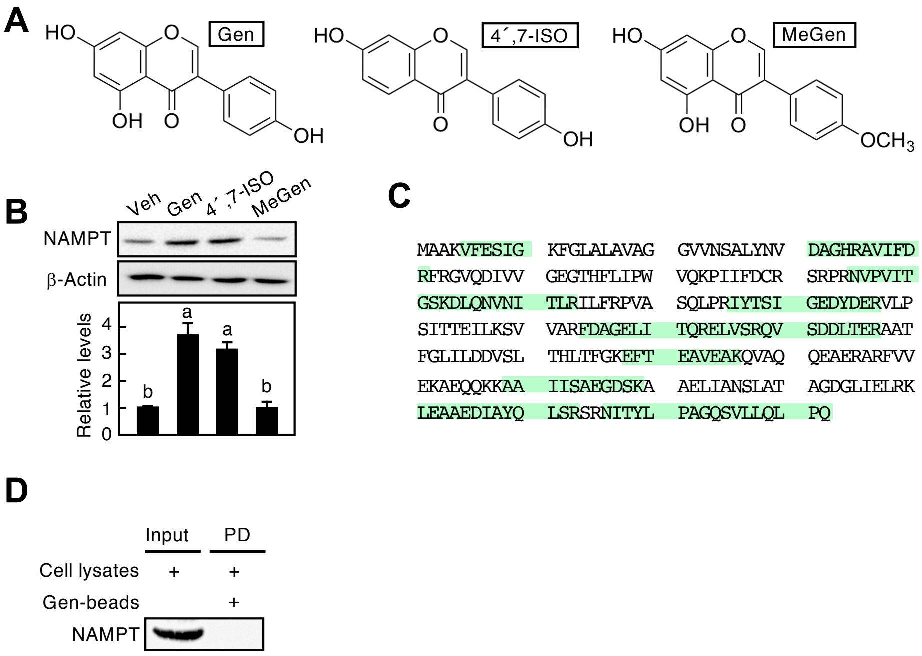


**Fig. S2. (**A) Chemical structure of isoflavones. (B) Western blotting of NAMPT in 3T3-L1 adipocytes after induction of differentiation with 10 μM isoflavones (genistein; Gen, 4′,7-dihydroxyisoflavone; 4′,7-ISO, 4′-methylgenistein; MeGen) for 7 days. (C) Amino acid sequences of PHB1. Obtained peptide fragments by nanoLC Q-TOF/MS and ProteinLynx Global server software are shown in green sheet. (D) Pulldown assay between Gen-bead and NAMPT. 3T3-L1 adipocyte lysates were incubated with Gen-bead, followed by western blotting with anti-NAMPT antibodies.
