## Supplementary Figure 3 for "Genistein enhances NAD^+^ biosynthesis by binding to Prohibitin 1 and upregulating nicotinamide phosphoribosyltransferase in adipocytes"

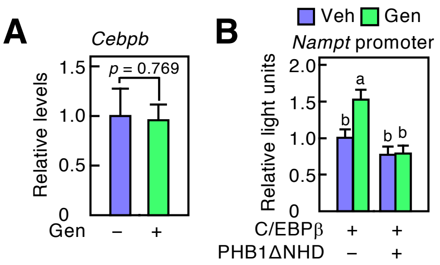


**Fig. S3.** (A) qPCR analysis of *Cebpb* in 3T3-L1 adipocytes after induction of differentiation with genistein (Gen; 10 μM). (B) *Nampt* promoter activity in 3T3-L1 adipocytes transiently transfected with pGL4-*Nampt*-Luc vector, and C/EBPβ and PHB1ΔNHD expression vectors, followed by incubation with Gen. Data are representative of triplicate independent experiments presented as the mean ± SD (n = 3). Statistically significant differences ($p < 0.05$) are indicated using different letters—a common letter between the groups indicated that the difference is not statistically significant.
