## Supplementary Figure 4 for "Genistein enhances NAD^+^ biosynthesis by binding to Prohibitin 1 and upregulating nicotinamide phosphoribosyltransferase in adipocytes"

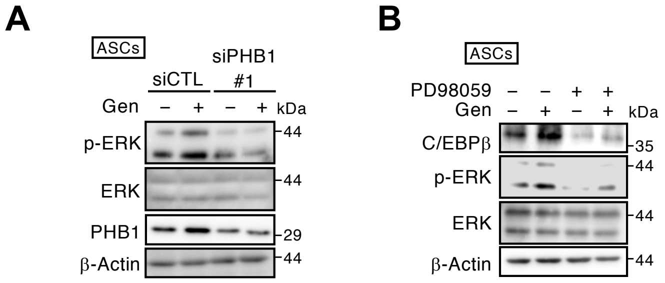


**Fig. S4.** (A) Phosphorylation levels of ERK in PHB1-specific siRNA (siPHB1)-treated ASCs treated with genistein (Gen; 10 µM). (B) Western blotting of C/EBPβ, p-ERK in ASCs treated with Gen and ERK inhibitor PD98059 (20 µM). All data shown are each representative of triplicate independent experiments.
