## Supplementary Figure 5 for "Genistein enhances NAD^+^ biosynthesis by binding to Prohibitin 1 and upregulating nicotinamide phosphoribosyltransferase in adipocytes"

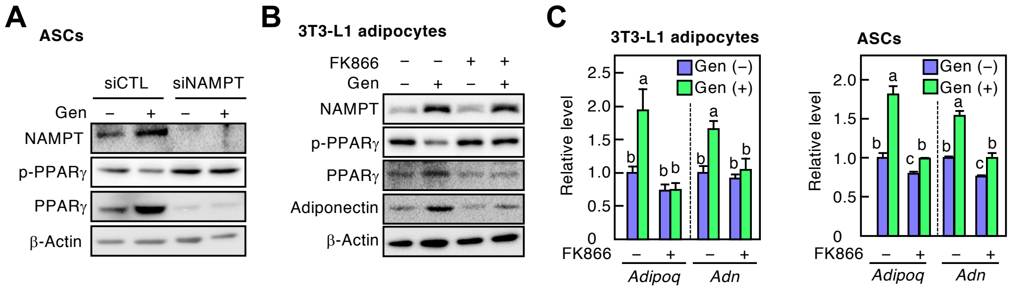


**Fig. S5.** (A) Western blotting in NAMPT-knocked down ASCs. The cells were transfected with siRNA for control (siCTL) or NAMPT specific (siNAMPT), followed by differentiation with genistein (Gen; 10 μM) for 7 days. (B) Protein expression of p-PPARγ, PPARγ and adiponectin in 3T3-L1 adipocytes. The cells were induced into adipocyte differentiation with Gen and/or FK866 (10 nM) for 7 days. (C) qPCR analysis in 3T3-L1 adipocytes (*left panel*) or ASCs (*right panel*) treated with FK866 in the presence or absence of Gen for 7 days. Data are representative of triplicate independent experiments presented as the mean ± SD (n = 3). Statistically significant differences ($p < 0.05$) are indicated using different letters—a common letter between the groups indicated that the difference is not statistically significant.
