## Supplementary Figure 6 for "Genistein enhances NAD^+^ biosynthesis by binding to Prohibitin 1 and upregulating nicotinamide phosphoribosyltransferase in adipocytes"

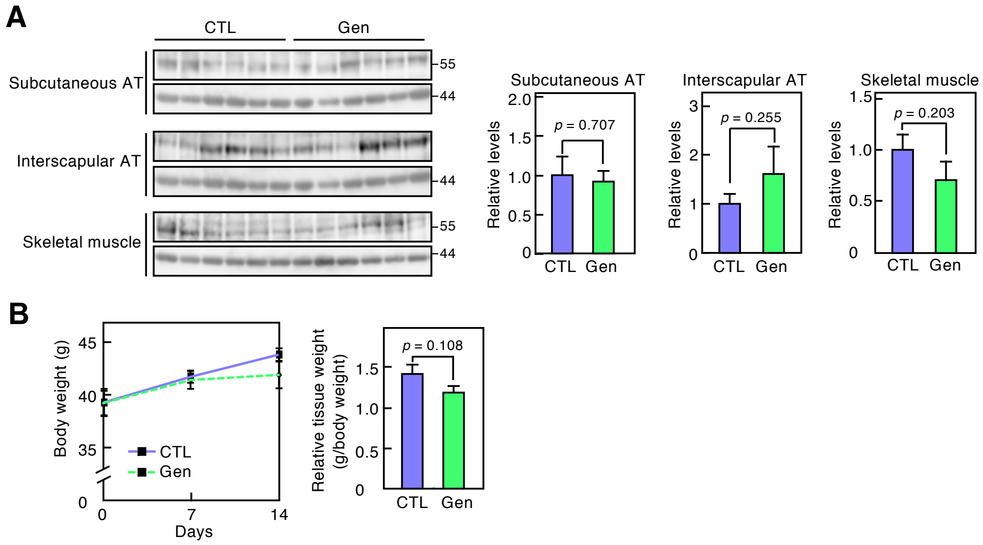


**Fig. S6.** (A) Western blotting of NAMPT in adipose tissues (AT) and skeletal muscle from mice following two weeks treatment with either vehicle (Veh) or genistein (Gen) (*left panels*). The ratio of each band was normalized to that of β-actin (*right* panels). (B) Body and visceral adipose tissue weight in Gen administered mice. Error bars represent the mean ± SEM (n = 6), and all data did not show statistically significant differences between the Veh and Gen groups ($p$ > 0.05).
